# A *Numerus* Population Viability and Harvesting Analyses Web App

**DOI:** 10.1101/074492

**Authors:** Wayne M. Getz, Oliver Muellerklein, Richard Salter, Colin J. Carlson, Andrew J. Lyons, Dana Seidel

## Abstract

Population viability analysis (PVA) is used to assess the probability that a biological population will persist for a specified period of time. Such models are typically cast as Markov processes that may include age, stage, sex and metapopulation structures, density-dependence and ecological interaction processes. They may also include harvesting, stocking, and thresholds that trigger interventions. Here we present Numerus PVA, which is a web app that includes extensible user-selected options. Specifically, Numerus PVA allows for the specification of one to ten age classes, one or two sexes, single population or metapopulation configurations with 2 or 3 subpopulations, as well as density-dependent settings for inducing region-specific carrying capacities. Movement among subpopulations can be influenced by age, metapopulation connectivity, and attractivity of regions based on the relative fitness of the youngest age classes in each region. Simulations can be carried out deterministically or stochastically, with a user-specified combination of demographic and environmental processes. Numerus PVA is freely available at http://www.numerusinc.com/webapps/pva for running directly on any browser and device. Numerus PVA is easily modified by users familiar with the NovaModeler Software Platform.

## Introduction

Highly valued animal and plant populations are declining at a dramatic rate worldwide (Barnosky et al. 2011), either as a result of overexploitation (Mullon et al. 2005, Godfray et al. 2010, Weinbaum et al. 2013), poaching (Chapron et al. 2008, Wittemyer 2011), global climate change, or the conversion of human settlement and development of pristine habitat (Ellis 2011, Foley et al. 2011, Urban 2015). The value of individuals within these populations comes from either ethical considerations (the conservation imperative: (Minteer and Collins 2010) or resource utility considerations (the management imperative: (Ostrom et al. 1999)). Since the seminal work of Beverton and Holt six decades ago (Beverton and Holt 1957), mathematical models have played a central role in both evaluating (Metzger et al. 2010) and devising population management programs (Weinbaum et al. 2013), initially for exploitation and, more recently, conservation. The latter has been implemented through creating protected areas (Moffitt et al. 2011), improving their security (Watson et al. 2014), and stocking or translocating individuals (Armstrong and Seddon 2008, Seddon et al. 2014) to bolster or reestablish populations in selected areas. At the core of the most comprehensive and successful of these models is the Leslie matrix formulation (e.g. see Caswell 2001), which provides a way of incorporating population vital rates (mortality and natality) into both harvesting (sustainable management) and population viability analysis (conservation management) models.

Impelled by the work of Beverton and Holt (Beverton and Holt 1957), fisheries science has had a cadre of quantitatively trained individuals able to formulate and code sophisticated models used to manage fisheries by helping set quotas on harvesting effort and fish stock removals (Getz and Haight 1989, Quinn and Deriso 1999). From the 1980’s onwards, quantitative population biologists have formulated Leslie matrix type models to help set quotas for trophy hunting or other types of exploitation of vertebrate populations (Getz and Haight 1989), but it is only over the past 20 years that the application of Leslie matrix type models has found wide application in conservation biology (Beverton and Holt 1957, Heppell 1998, Menges 2000, Wisdom et al. 2000, Caswell 2001, Fieberg and Ellner 2001, Crone et al. 2011, Merow et al. 2014b). In some cases, when traits, such as age or size are considered as continuous rather than discrete variables, these models are more generally formulated as integral projections (e.g. see Easterling et al. 2000, Merow et al. 2014a); though they revert to matrix models under numerical discretization schemes (Ellner and Rees 2006, Rees et al. 2014). With a rapidly growing need to conserve endangered species, scientists and policy markers who have not been trained to code their own population models for numerical simulation, face the challenge of building best practice simulation models (Kettenring et al. 2006) to aid them in their species management or conservation biology work. To support these researchers and managers, software applications platforms, such as RAMAS (e.g. see Crone et al. 2011) and VORTEX (Lacy 1993, 2000, Brito and Da Fonseca 2006, Lacy and Pollak 2012), have been developed, particularly to aid population ecologists in using population viability analyses (PVA) (Beissinger and Westphal 1998, Morris and Doak 2002). VORTEX (Lacy and Pollak 2012) takes an agent-based approach to modeling individuals and is able to include the type of information used to track lineages and pedigrees that are stored in studbook files (cf. Marker and Fund 2012). VORTEX is also able to track, under the assumption of Mendelian inheritance, the fate of multiple genetic loci and assess levels of inbreeding. VORTEX runs in a Windows operating systems environment, as does RAMAS. RAMAS, though, is a commercial platform with multicomponent high-end products that can incorporate detailed landscape information and geographical information systems approaches into its analyses.

Our *NumerusOL* app, which we refer to as Numerus PVA, provides a more gentle entry into PVA than either VORTEX or RAMAS. Most importantly, Numerus PVA runs directly in a web-browser environment with access via the website Numerusinc.com. Further, it has sufficient flexibility to include two sexes, a variable sex ratio, three types of density-dependent mechanisms, and runs as either a deterministic or stochastic simulation (demographic, environmental or both) with one to three subpopulations. Migration of individuals among subpopulations incorporates propensity of individuals to move by age and sex, connectivity of regions, and attractivity of regions based on some criterion such as the anticipated fitness of individuals within those regions. Harvesting and stocking management options are also included. All of these components are wrapped in an intuitive web-based Graphical User Interface (GUI) that requires no computer programming and only an elementary understanding of discrete time population models using life table data.

Numerus PVA itself was constructed using the Numerus platform and the *NumerusOL* file conversion technology, which is based on our earlier Nova Platform (Salter 2013, Getz et al. 2015). Use of Numerus PVA, however, requires no knowledge of coding nor of the Numerus modeling platform. The GUI provides data fields that can be entered online and key parameters manipulated using sliders. The app can be run in either deterministic or stochastic modes. The deterministic mode is most useful when evaluating various management strategies implemented in large population (generally in the context of sustainable fisheries or forestry exploitation rates that are optimal in some sense—cf. Getz and Haight 1989). The stochastic mode includes demographic stochasticity (e.g. small population size effects) and environmental stochasticity (e.g. driving variable fluctuations drawn from climatic variable distributions). The former is critical to carrying out species extinction risk analyses, in other words PVA, (Fieberg and Ellner 2001), while the latter permits possible climate trend information to be accounted for in multi-decadal simulations (cf. Wilmers and Getz 2004a).

## Model Structure and Simulation Modes

### Demography

The demographic model underlying our app has the flexibility to include one or two sexes, single or metapopulation structure, and density-dependent (DD) effects (Fig. 1). These DD effects can be implemented separately in each metapopulation region and in the context of reducing survival rates in the youngest male and female, oldest male and female, and maturing male age-classes. In all but the latter, we assume that survival is affected by the aggregated biomass variable *B* (Fig. 1) that weights individuals by a user-specified age-sex relative value, rather than purely by the more usual population size (i.e. numbers). This allows us to correct for the fact the when density dependence arises through resource consumption then total biomass is more appropriate then population size for indexing the level of competition (Getz 2011). In contrast, maturing male DD effects depend on the total number of males *M* that are ≥ age *r* (age at which males mature; see figure in Appendix 4, SI online) because we assume these effects arise through contest competition or defense of territories. In essence, the model depicted in Fig. 1 is a nonlinear elaboration of a discrete-time Leslie matrix formulation (Caswell 2001), with details of the equations provided in Appendix 1 (SI).

**Figure 1.**
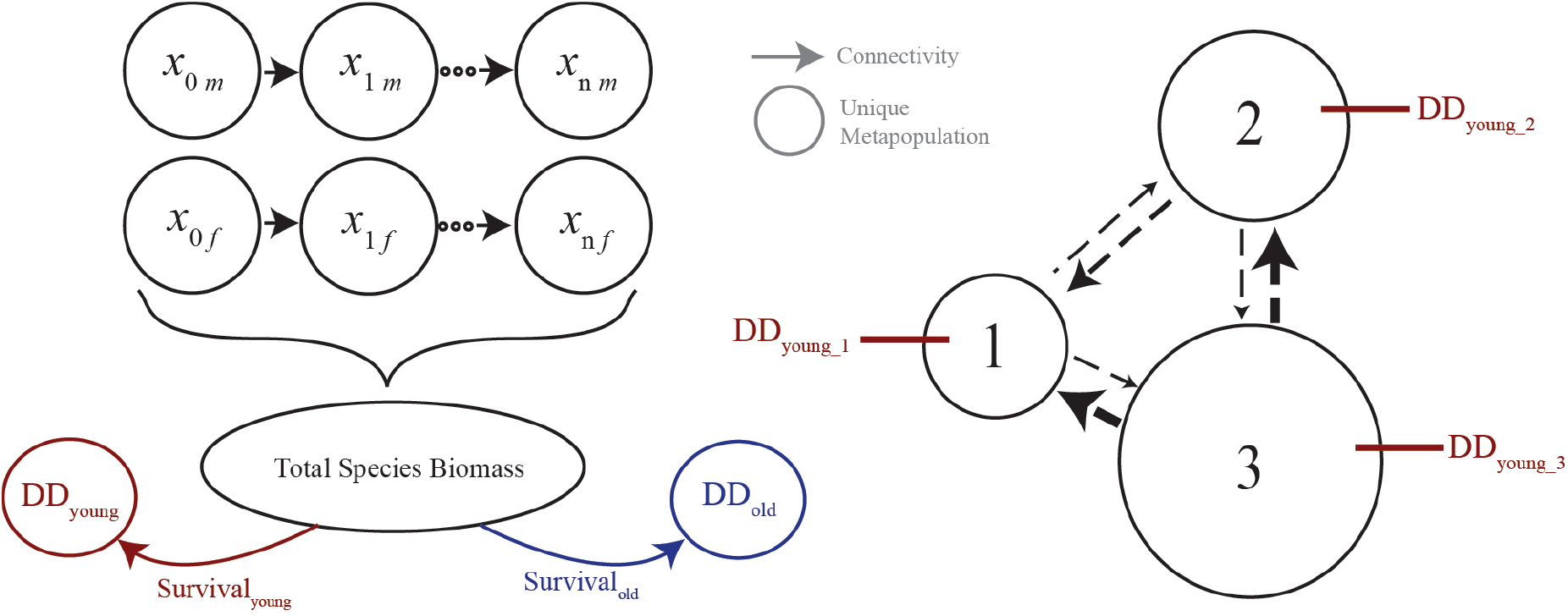
A flow diagram of a two-sex, age-structured, population model with density-dependent (DD) survival in the youngest and oldest classes that depend on the population biomass *B* (Eq. 3) (left side), and schematic of the metapopulation structure (right side) indicating regionally specific DD processes.

The data needed to implement a Leslie matrix model of a homogeneous population are the age-specific survival rates *s*_*i*_ (the proportion individuals that survive from age *i* to age *i*+1) and the age-specific natality values *b*_*i*_ (the average number of newborns per unit time produced by adults of age *i*; or females-per-female when two sexes are invoked). If *x*_*i*_(*t*) is the number of individuals at time *t* in a model that does not differentiate by sex, then the number of newborns at time *t*+1 is (in Fig. 1 we illustrate the more general sex differentiated model with variables *x*_*if*_ and *x*_*im*_, *i*=0,…,*n*)

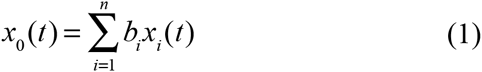
Aging is included in the model through the equations

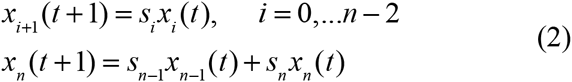
Density dependence in the model is introduced by multiplying either or both of the density independent survival rates *s*_0_ and *s*_*n*_ by the factor

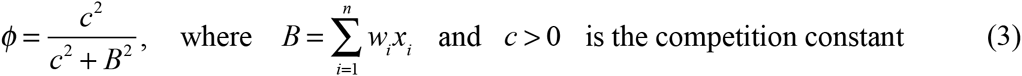
This dependence is then generalized to be included in two sex models; and it can also be applied to males that may be motivated to engage in male-male competition on reaching sexual maturity, as illustrated in Fig. 1. In this case the density dependent factor is

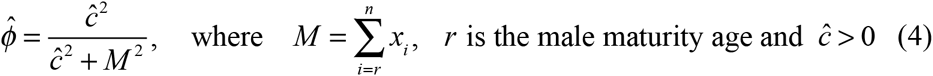
In addition, in a two-sex model, if there are no mature males during an iteration period, then births are set to zero for that period.

### Migration

To specify how individuals move among meta-population regions, a region-specific propensity-to-move vector **q**_*i*_ of elements *q*_*ij*_ is entered (Fig. 2): in particular, *q*_*ij*_=0.*x*, implies that individuals of age-sex class *i* in region *j* have an *x*% propensity to move. In addition a connectivity matrix with elements *p*_*rs*_ (Fig. 2) specifies the relative ease that individuals in region *r* are able to move to region *s*, should they have the propensity to do so. Thus the movement of individuals is determined both by this connectivity matrix and by a propensity for individuals of different ages to move at different rates (e.g. only maturing males in search of territory may move, and so on).

**Figure 2.**
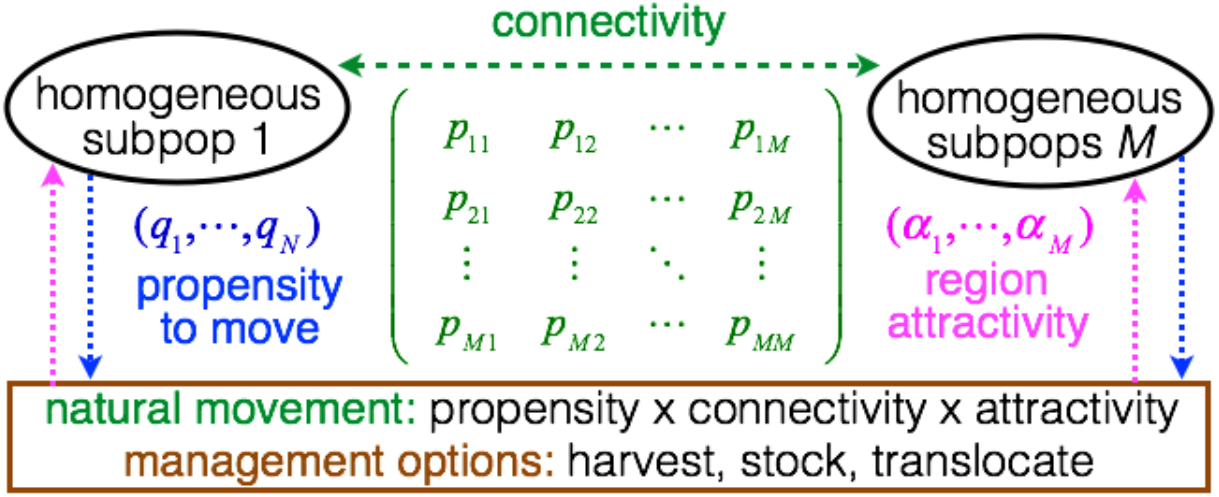
Depiction of the stochastic movement (i.e. migration) and management processes used to compute exchanges among and changes *M* ≤ 3 regional subpopulation levels, as generated by *harvesting* and *stocking* (which can also be interpreted as translocations) and then *migration* (cf. Fig. 3). The latter is computed in terms of i) a *propensity* of individuals of different age-sex classes to leave their current regional location, ii) a matrix that represents the *connectivity* (ease of movement) among regions; and iii) an optional destination *attractivity* factor determined by the relative fitness of the youngest age class in region (DD1 and DD2 in Fig. 1, as influence by *f* defined in Eq. 3).

The movement (migration) module of Numerus PVA has one additional feature referred to as the *engage-relative-fitness* switch, which can be turned ON or OFF. If it is OFF, the probability of individuals moving from one region to another is determined solely by the vectors ***q***_*i*_ and matrix ***P*** (Fig. 2). If this switch is on then an additional relative attractiveness factor is applied for each region: the value of these factors across regions are in proportion to amount that survival of the youngest age classes is reduced in each region by density-dependence (see Appendix 1, SI, for details).

### Stochasticity

The model can run either deterministically or stochastically, where the latter may include either demographic or environmental sources (Engen et al. 2005). Demographic stochasticity arises from sampling theory (underlying vital parameters are treated probabilistically) and hence should be included when populations are small, because its effects are proportional to the inverse of the square root of population size (Desharnais et al. 2006). Environmental stochasticity is likely to affect various age classes differently, depending on the source of the stochasticity (e.g. climatic drivers versus disease). Here we make provision for environmental stochasticity to impact only the youngest age class, which is often the most vulnerable age class, as has been documented for large mammalian herbivores (Gaillard et al. 1998). Environmental stochasticity may also be important in senescing age classes (Wilmers and Getz 2004b). Such considerations can be included in customized versions of the model, as discussed in the Conclusion Section. A deterministic simulation requires the demographic stochasticity switch to be OFF, and the environmental stochasticity slider to be set to 0. If demographic stochasticity is ON, then all survival computations of the form *s*_*i*_*x*_*i*_ are replaced with BINOMIAL(*x*_*i*_, *s*_*i*_) computations and birth number computations *b*_*i*_*x*_*i*_ are replaced with BINOMIAL(*b*_*i*_*x*_*i*_,*b*_max_/*b*_*i*_) computations, where *b*_*i*_(birth rate) has the interpretation of expected litter size and *b*_max_ (birth max) is the maximum litter size. If the environmental stochasticity slider is > 0 then environmental stochasticity in the survival *s*_0_ of the youngest age class is included up to a maximum level that is implement when the slider is set to 1 (see Appendix 1, SI, for more details). The slider value can be tuned over multiple simulations with different slider values until the simulated variance matches the desired or observed variance.

### Constraints

The current version of the software is limited to 1–10 age-classes for each of 1–2 sexes in each of 1–3 metapopulation regions and running the model for a maximum of 1000 steps. Future versions of the software will relax these constraints. The NovaScript file, underlying this *NumerusOL* implementation, was constructed using the Numerus modeling platform (Salter 2013, Getz et al. 2015) and it (see SI) can always be rapidly modified to supply the user with a version of the model that meets the user’s needs.

### Event Sequence

Since the order of events matters in a discrete time simulation—e.g. an individual that dies first cannot then be harvested, and vice-versa—it is necessary to pay attention to this order when formulating transition equations from one time step to the next. The order that we use to calculate the number of individuals in age class *i* at time *t*, i.e. *x*_*i*_*(t)*, that make it into age class *i*+1 at time *t*+1, depends first on the number that die from natural causes (as determined by the survival parameter *s*_*i*_), then on those removed by harvesting, then on those added by stocking, and finally the numbers lost and gained through migration (Fig. 3). The implication of this is that individuals that move do not include those that are harvested, but may include those that are stocked. Also, the density-dependent mortality depends on the population state at the beginning of the discrete interval of time and hence does not account for population changes from harvesting, stocking, or movement.

**Figure 3.**
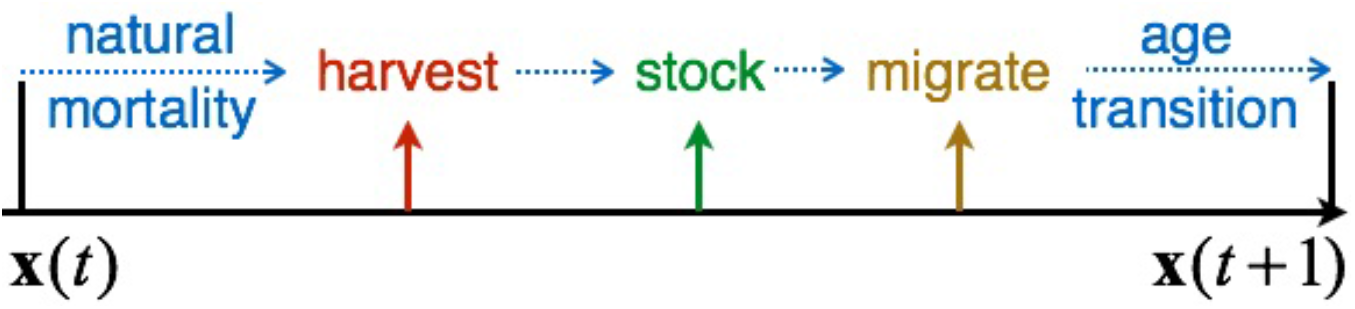
Sequence of events used to compute the transition of the age class vector **x** from times *t* to *t*+1. From this we see that density-dependent (DD#, cf. Fig. 1) survivorship, when applied, takes effect prior to harvesting, stocking and migration.

## Setting up a Model Run

At the simplest level, one can choose to run the model in deterministic mode for a single-sex homogeneous population with no density dependence. At its most complex, the Numerus PVA can be run in two-sex, density-dependent, metapopulation mode, with both demographic and environmental stochasticity switched on, where all parameters are metapopulation region specific. The values of parameters can either be entered manually or by importing an appropriately configured csv file (Fig. S1, SI online). Using the manual entry approach (with or without the self-guided tutorial) the data that are entered on the following pages, which appear sequentially:

i. *Population data*: number of regions; number of age classes, one or two sex, and male-maturity-age by region in the two-sex case, density dependence options by region (young, old, or mature males in the case of two-sex case) (Fig. S2, SI online). These data are used to set up the forms for the pages that follow, since the much of the remaining data is age-class and region specific.
ii. *Core population parameters*: initial numbers by age for each region, survival and birth rates by age for each region, “birth max” by age (will only be used for models with demographic stochasticity) for each region, and finally both female sex ratio and male maturity age by region. These data pertain to the entries needed to implement a Leslie Matrix population projection model.
iii. *Biomass and movement parameters*: relative biomass of each of the age/sex classes in each region; density-dependence parameters (youngest age class, oldest age class, males transitioning to sexual maturity) in each region; propensity to move by age/sex class for each region; region connectivity matrix. These data are used to implement the density dependent functions that modify survival rates of the youngest, oldest, and maturing male age classes in each region, as well as determine stochastic movement rates among regions.
iv. *Stocking and harvesting values*: age-class specific stocking and harvesting rates (numbers to be added or removed) each time interval or on a regular schedule with adjustable frequencies; and additional harvesting pressure that removes specified numbers of individuals, but randomly from cohort ranges set by sliders.
v. *Interactive model implementation*: a page from which runs are executed once with the option to control the following settings or switches (Fig S3, SI online): simulation length setting, save model output switch; pseudoextinction threshold setting, density dependent engagement switches; demographic stochasticity switch, environmental stochastic switches and levels by region, as well as random harvesting values (number specified, by individuals chosen from random classes within specified cohort ranges). On running the model visual output will be generated (Fig. S4, SI online) and an optional csv file generated.

## Illustrative Example

We illustrate implementation of our PVA web app, using an exemplar data set that is inspired by the life-history and conservation predicament of the black rhino, *Diceros bicornis*, in southern Africa; but has not been fitted to any particular population because parameters vary quite considerable among populations. This species, like other rhino species, is close to extinction (it is on the IUCN’s critically endangered list—next step, extinct in the wild), with fewer than a few thousand individuals alive at this time. Because this species is subject to the devastating effects of intense poaching for rhino horn, actual locations and numbers are kept confidential by managers of national parks and conservation areas. Further, while some life history data on birth and survival rates are available, these rates vary from one area to another, and often life table construction (natality and mortality rates at each age) relies on misleading values obtained from individuals kept in zoos (e.g. longevity in zoos can be greatly different from longevity in the wild; while calf survival depends on predation pressure). Thus we stress that our dataset exemplar should not be regarded as applicable to any specific rhino population and the model itself is essentially generic. The analysis that follows here is not meant to apply to any real population, but is provided for the purpose of illustrating how conservation decisions for the species can be evaluated using our Numerus PVA app.

### Basic population parameters

The inter-calf interval of mature female rhinos is approximately 3+ years (includes 1.3 years for gestation). For this reason, it is convenient to organize the population into age classes that each span three years: i.e., the basic iteration units for *t* in the model will be 3-year intervals. Hence if the model is used to project population change over *T* units of time, the corresponding number of years for the projection is 3*T* (Note: in Figs. 4 and 5 the *x*-axis denotes units of *t*, while in the Fig. 6 the units are years rather then *t*). Recent estimates of calf, adult female, and adult male survival of rhino in an area of Namibia regarded as relatively unproductive for rhino growth was respectively 0.793, 0.944 and 0.910 (Brodie et al. 2011). These data, rounded to the nearest 0.05, are listed in Table 1, along with the estimate that females in this region produce 0.315 calves per female per year. The female sex ratio used is 0.6, based on estimates reported in Law et al. 2014, and the male maturity age is taken to be 2 (age 6^+^-9). These data, when used in a 9-age class, female only Leslie matrix model yield a density-independent growth rate of 6.4% per annum (see SI Appendix 3).

**Table 1.** Life Table of a Large Mammalian Herbivore

| Age (class; $i$) | Male survival | Female survival | Young / female | Female weights* | Male weights* |
| --- | --- | --- | --- | --- | --- |
| 0-3 (calf: 1) | 0.80 | 0.80 | 0 | 0.5 | 0.5 |
| 3<sup>+</sup>-6 (subadult: 2) | 0.90 | 0.95 | 0.3 | 0.75 | 1 |
| 6<sup>+</sup>-9 (young adult: 3) | 0.90 | 0.95 | 0.95 | 0.75 | 1 |
| 9<sup>+</sup>-12 (mature adult: 4) | 0.90 | 0.95 | 0.95 | 0.75 | 1 |
| 12<sup>+</sup>-15 (mature adult: 5) | 0.90 | 0.95 | 0.95 | 0.75 | 1 |
| 15<sup>+</sup>-18 (mature adult: 6) | 0.90 | 0.95 | 0.95 | 0.75 | 1 |
| 18<sup>+</sup>-21 (mature adult: 7) | 0.90 | 0.95 | 0.95 | 0.75 | 1 |
| 21<sup>+</sup>-24 (mature adult: 8) | 0.90 | 0.95 | 0.95 | 0.75 | 1 |
| >24 (aging adult: 9) | 0.60 | 0.60 | 0.5 | 0.75 | 1 |

\*relative units in terms of resource consumption

### Density dependence

The carrying capacity for rhino in two Zululand parks in South Africa, has been estimated at 0.4 and 1.6 rhino km^−2^ (Conway and Goodman 1989), while values for this species in drier regions of southern Africa have been estimated to be as low as < 0.1 rhino km^−2^ (Linklater et al. 2011). In this illustrative example, we include the effects of density-dependence only on the survival of the young: that is DD1 and DD2 in Fig. 1, which is tantamount to multiplying the density independent survival constants for males and females in Table 1 by the function given in Eqn. 1. We note that setting *c*=50, 100, and 150 producing carrying capacities (i.e. equilibrium values) of around 69, 138 and 208 individuals (Fig. S6; implying reserves of corresponding sizes in square kilometers).

### Stochasticity

For purposes of comparison, we run the model with the parameter values specified in Table 1, the sex ratio at 0.6 (female biased), and *c*=100 in deterministic, and environmental stochasticitiy at half max and at full max settings (the environmental stochasticity sliders are set at 0, 0.5 and 1 respectively) (Fig 4A). We repeated this simulation with demographic stochasticity switched ON (Fig. 4B). First we note from visual inspection of Fig. 4 that variance increases with increasing levels of environmental stochasticity and that the model predicts whole numbers when demographic stochasiticity is ON, but fractional numbers when demographic stochasticity is OFF. This is consistent with the requirement that demographic stochasticity be ON when population size is relatively small (tens or hundreds) and can only be safely ignored when population sizes are close to a thousand or more, in which case the interpretation of the state variables is density (i.e. fractional numbers are meanful) rather than size (fractional numbers are nonsensical).

**Figure 4.**
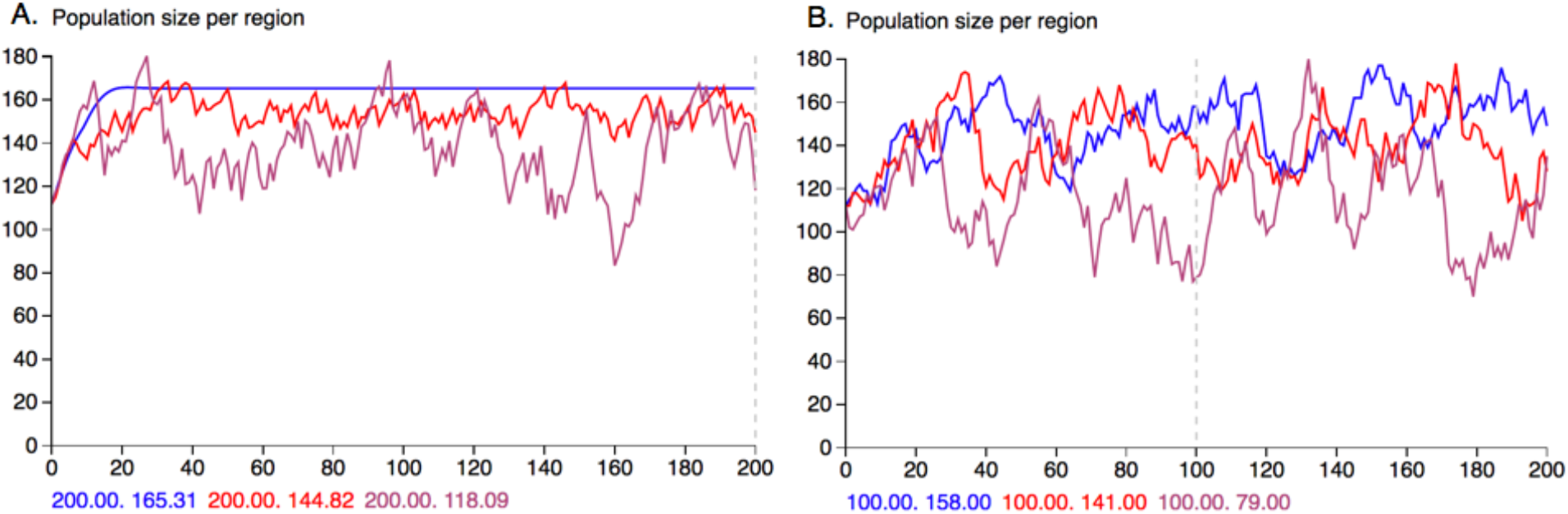
Simulations of model using parameters in Table 1, female sex-ratio=0.6 and male maturity age is 3 in all three regions, and density-dependent survival is only in the youngest group (parameter *c*100), with no (blue), half (red) and maximum (purple) levels of environmental stochasticity with demographic stochasticity OFF (A.) and ON (B.). The size of the population is read out *t*=200 (A.) and *t*=100 (B.) where we note that when demographic stochasticity is OFF (A.), the model predicts fractional numbers, while when demographic stochasiticity is ON, the model predicts whole numbers. Initial values for male and female cohorts in all cases were: male=(10,9,8,7,6,5,4,3,4)´ (´ denotes the vector is transposed from a column to a row) and female=(10,9,8,7,6,5,4,3,4)´)

### Movement

For the three components of migration—propensity to move, connectivity, and region attractivity (Fig. 2)—we allowed individuals only in the third age class to move (i.e. those aged 7–9 years, through the propensity to move vectors: male=(0,0,1,0,0,0,0,0,0)´; female=(0,0,1,0,0,0,0,0,0)´), we assumed all regions where equally accessible for another (i.e. the connectivity matrix was filled with 1’s); and we assumed all regions were equally attractive (i.e. the “Migration with Relative Fitness” switch was OFF). Notice that extinction occurs in region 3 without migration (Fig. 5A.), but the populations in the three regions are somewhat equalized when migration occurs (Fig. 5B.).

**Figure 5.**
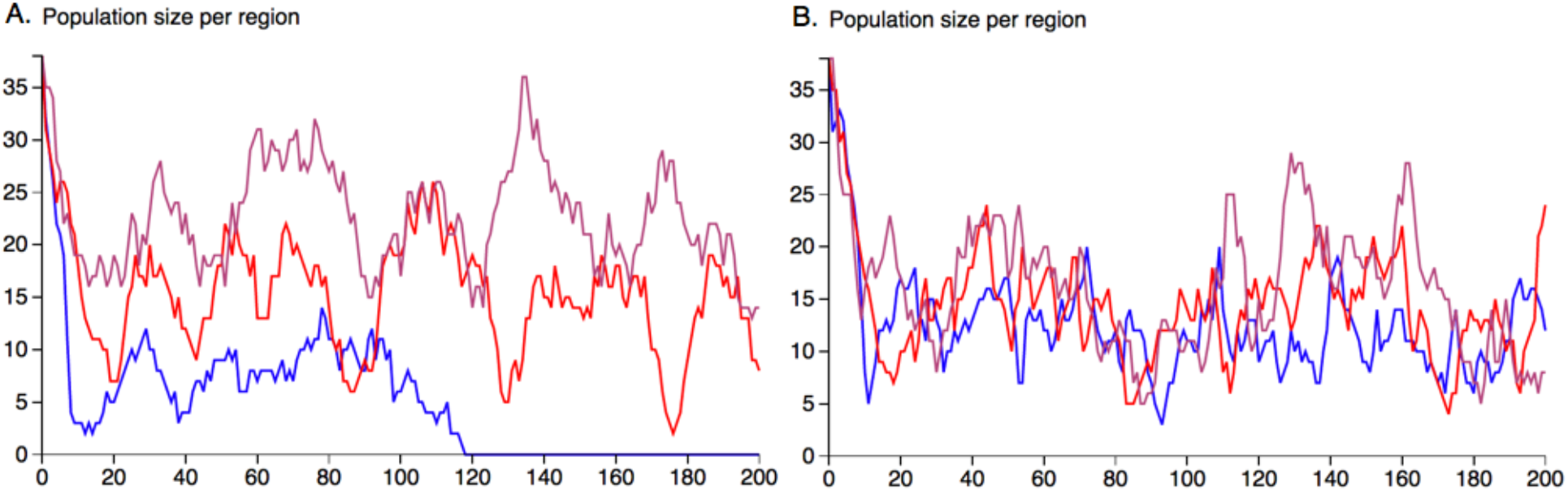
Screen captured simulation output from Numerus PVA using parameters in Table 1, sex ratio = 0.6, and density-dependent survival in the youngest age group in each of three regions, where the parameter *c* has values *c*=5 (blue), *c*=10 (red) and *c*=15 (purple), respectively. The cases of no migration among regions (A.) and movement of males and females in age class 3 (i.e. 7–9 year olds) only among regions (B.) are illustrated. Initial values for male and female cohorts in all cases were: male=(3,3,3,3,2,2,1,1,1)´ and female=(3,3,3,3,2,2,1,1,1)´.

### Poaching

We carried out an assessment of the effects of poaching by running the model with parameters used to generate Fig. 4, but with youngest age group density-dependent survival parameter set at *c*=10 (i.e. the red trajectory in Fig. 5A applies) under both *no poaching* (harvesting) and *poaching* scenarios. We obtained 36 replicate runs for the no poaching scenario by running the model in 3-region mode 12 times under the assumption of no migration, where the red trajectory in Fig. 5A is but one example. We then reran with harvesting set to removing 1 male and 1 female in each time period drawing the individual at random from cohorts 4–9 for the males and 5–9 for the females. This level of poaching considerably increases the population’s risk of extinction from comfortably less than 10% over a 99-year interval (simulation interval is 33 time units) (blue curve, Fig. 6) to almost 90% (red curve, Fig. 6)

**Figure 6.**
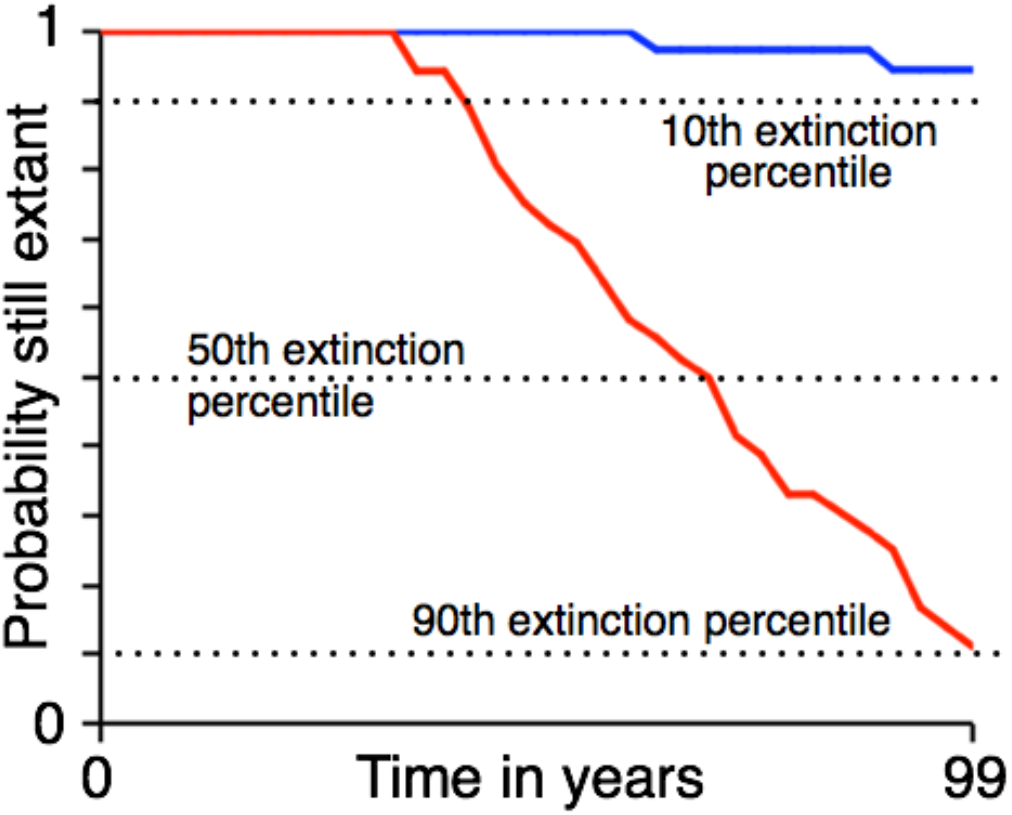
Extinction curves obtained for data generated by Numerus PVA for no harvesting (blue) and random harvesting (red; see text for details) of 1 male and 1 female in each time period. Data obtained from 36 simulations (i.e. 12 repeated simulations in 3-region mode with no migration) in each case.

## Conclusion

The Numerus PVA app presented here when in its one region, one sex, no density dependence, deterministic mode can be used as a class room tool to introduce students to the behavior of Leslie matrix (discrete time, linear, age-structured) models. The app can also be used to introduce students to discrete stochastic process models and their application to PVA risk analysis by enabling demographic stochasticity (i.e. flipping its switch on the app) and incorporating density-dependence into the youngest or oldest age class. Additionally, the app can also be used in the classroom to introduce notions of population management through both harvesting and stocking actions, as well as for exploring migration processes in metapopulation settings. Beyond this the Numerus PVA app can be use in research, though the current constraints limit its application to 20 age-sex classes and 3 regions. These constraints are easily relaxed and users can contact Numerus (at Numerusinc.com) to obtain more powerful versions and make extensive runs on high performance computing clusters.

## Glossary

*Biomass:*: The total mass of a population is commonly used in ecology and resource management in lieu of population size as an alternative to the number of individuals. In Numerus PVA, biomass is used to implement density-dependence (DD) effects.
*DD effects:*: Demographic or environmental limits that reduce population growth as populations get larger. Numerus PVA provides density-dependence options that affect survivorship of the youngest age classes youngest female (DD1) and male (DD2), oldest female (DD3) and male (DD4), and mature male (DD5) age-classes, as illustrated in Fig. 1.
*Demographic stochasticity:*: Random fluctuations arising from the probabilistic nature of applying vital rates to individuals at every life stage in both sexes.
*Environmental stochasticity:*: Random, environmentally-induced fluctuations in survivorship. In Numerus PVA, environmental stochasticity is an option for survivorship of the first life stage only (i.e. environmentally-induced juvenile mortality).
*Leslie matrix:*: A transition matrix underlying a discrete-time, linear, age-structured population dynamic model.
*Metapopulation:*: A set of connected subpopulations. In Numerus PVA, metapopulations are modeled as a weighted node network with implicit movement of individuals along vertices.
*Metapopulation connectivity:*: An underlying matrix with entries, scaled to take values on the interval [0,1], that represent the relative ease-of-transition among different nodes in the metapopulation.
*Perron root:*: The dominant eigenvalue of a square non-negative matrix; the Perron root of a Leslie matrix is the rate of population growth.
*Propensity to move:*: In Numerus PVA, movement propensity is an age- and sex-based demographic state specifying the likelihood of emigration to another area (independent of destination).
*Pseudoextinction:*: The event horizon of population size, below which extinction is certain. In Numerus PVA, pseudoextinction levels can also be treated as thresholds for interventions such as *ex situ* captive breeding programs.
*Regional attractivity:*: In Numerus PVA, once the decision to move has been made, and the connectivity of nodes accounted for, an intrinsic variability in quality of possible destination regions remains. We use the comparative intensity of density-dependent (DD) effects on survivorship of the youngest age class (quantity *φ* in Eq. 3, with *c* pertaining the youngest female age class) to scale this quality so that individuals are more likely to go to regions with smaller rather than bigger DD1 effects.

## Authors’ Contributions

The model was formulated by WMG, the application produced by OM, using technologies pioneered by RS. Training videos were created by AJL. All authors contributed to the look of the app. The draft was produced by WMG and all authors helped with the figures and editing of the text.

## Access Numerus PVA App

This web app can be accessed at www.numerusinc.com/webapps/pva/

## Supplementary Information (SI) online

## Appendix 1. App Data Entry

**Fig. S1.**
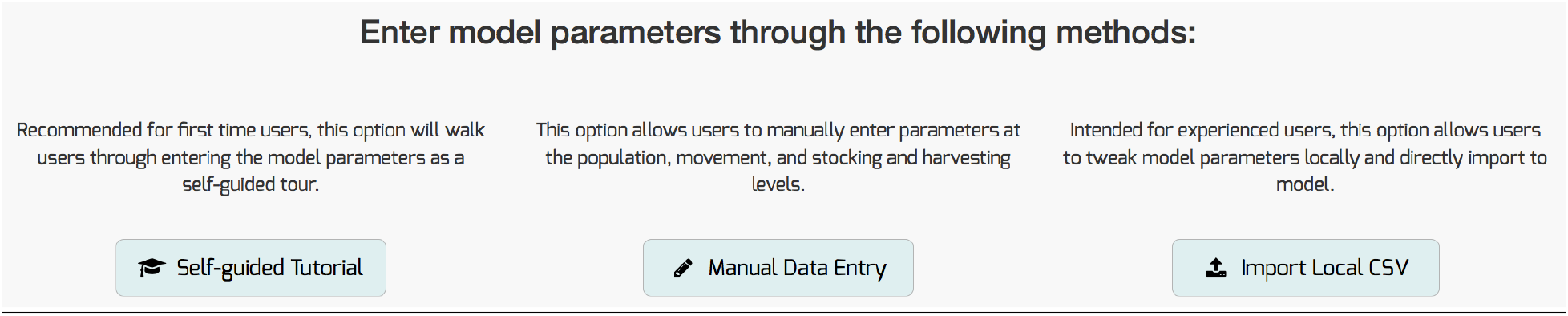
Data can be entered either online or by importing a local csv (comma separated values) file.

**Fig. S2.**
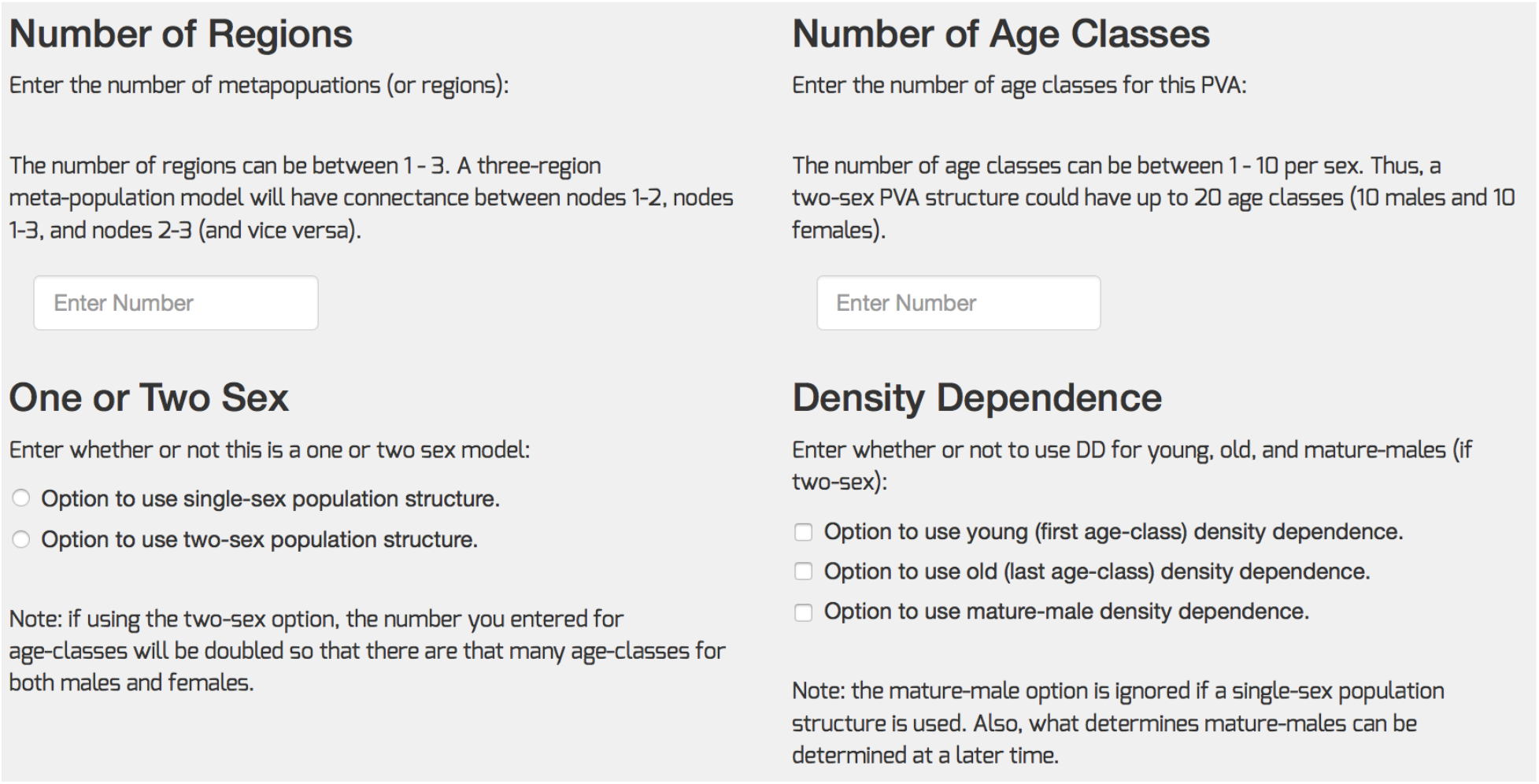
Population “core structure” data entry page.

**Fig. S3.**
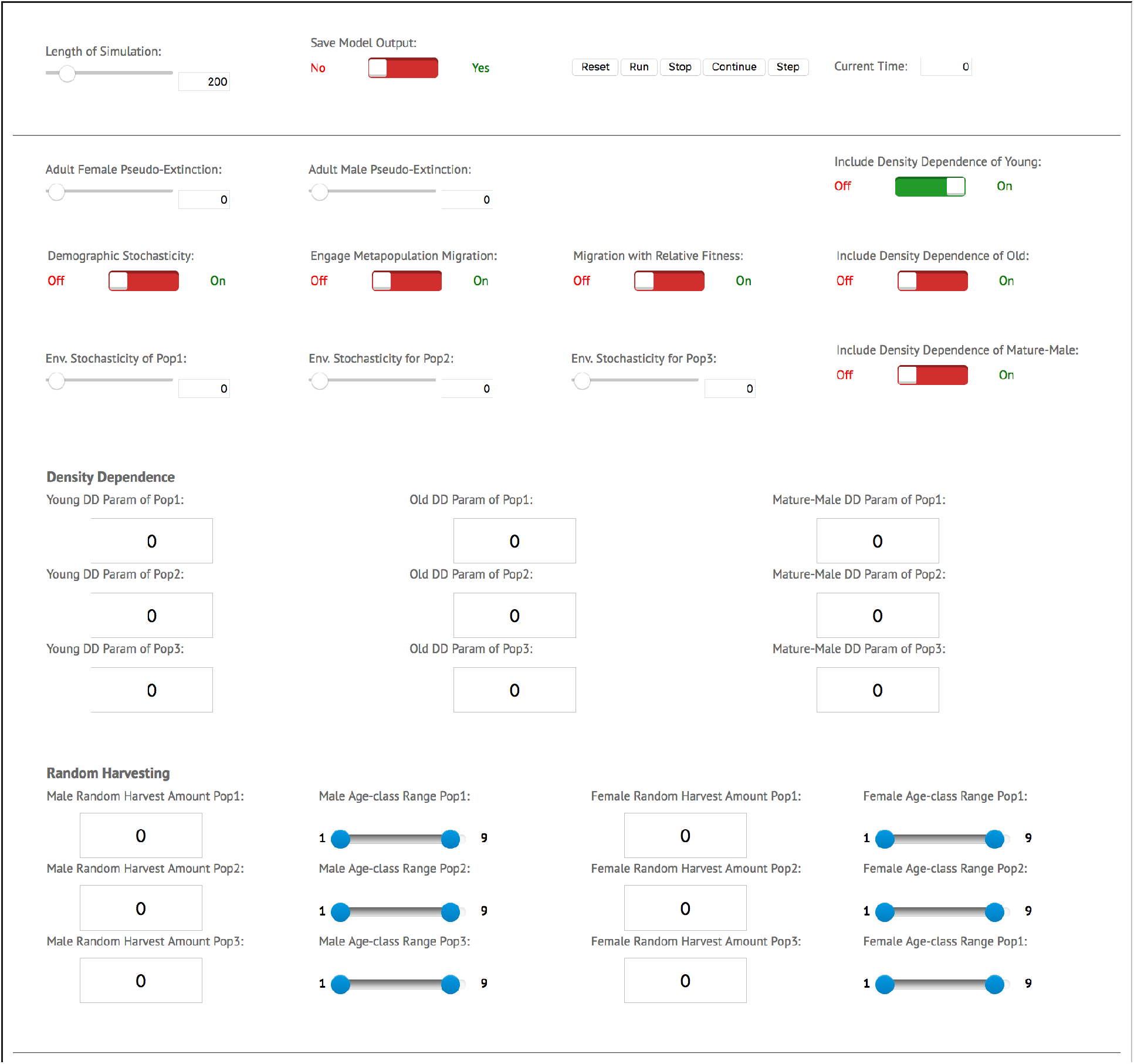
Model simulation control window.

**Fig. S4.**
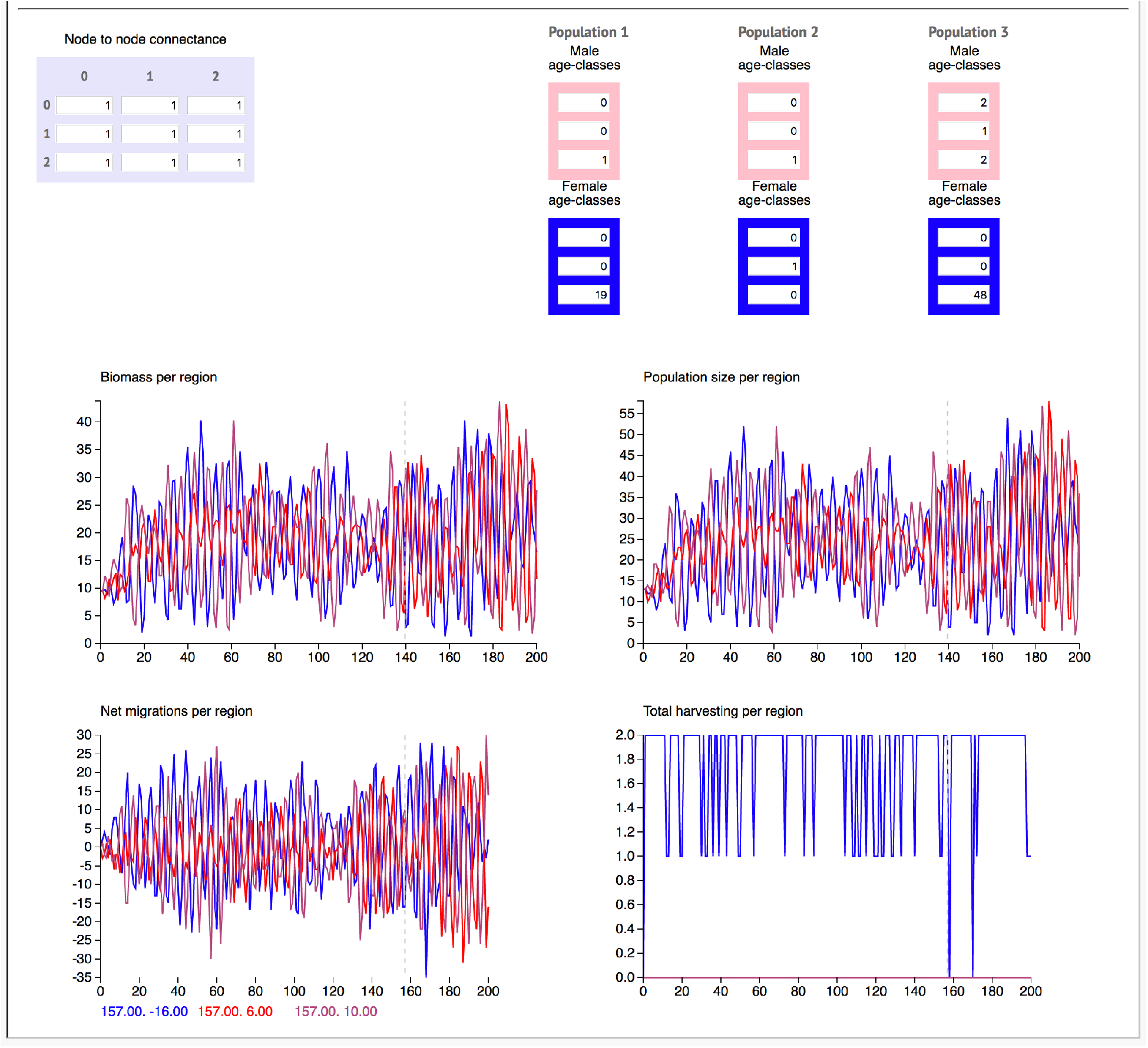
Output window.

## Appendix 2. Generic Rhino Example

The csv file used is available at the web app site www.numerusinc.com/webapps/pva/

### Equivalent Leslie Matrix model

Leslie matrix model constructed from Table 1 (terms in first row are births by age class multiplied by first period survival value 0.8 and sex ratio 0.6).

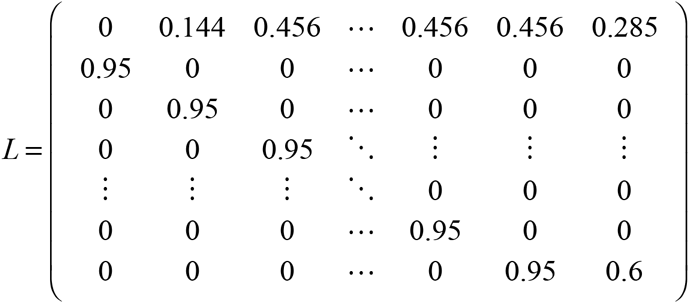
The dominant eigenvalue (Perron root) of this matrix is 1.20451. Since each time step is equivalent to three years, this represents a 20.5% increases per time period; i.e., a 6.4% per annum growth rate.

If we assumed the under sustained poor environmental conditions that *s*_0*l*_, *l*=*m*, *f*, drops precipitously from 0.8 to 0.2, then, if this were sustained in perpetuity, and eigenvalue analysis indicates that the population would decline at an annual rate of just over 2% per annum. If, however, the value of *s*_0*l*_, *l*=*m*, *f*, average out to be (0.8+0.2)/2=0.5 then eigenvalue of this average matrix yields a growth rate of 3.3% per annum.

### Density dependence of youngest age class

**Fig. S5.**
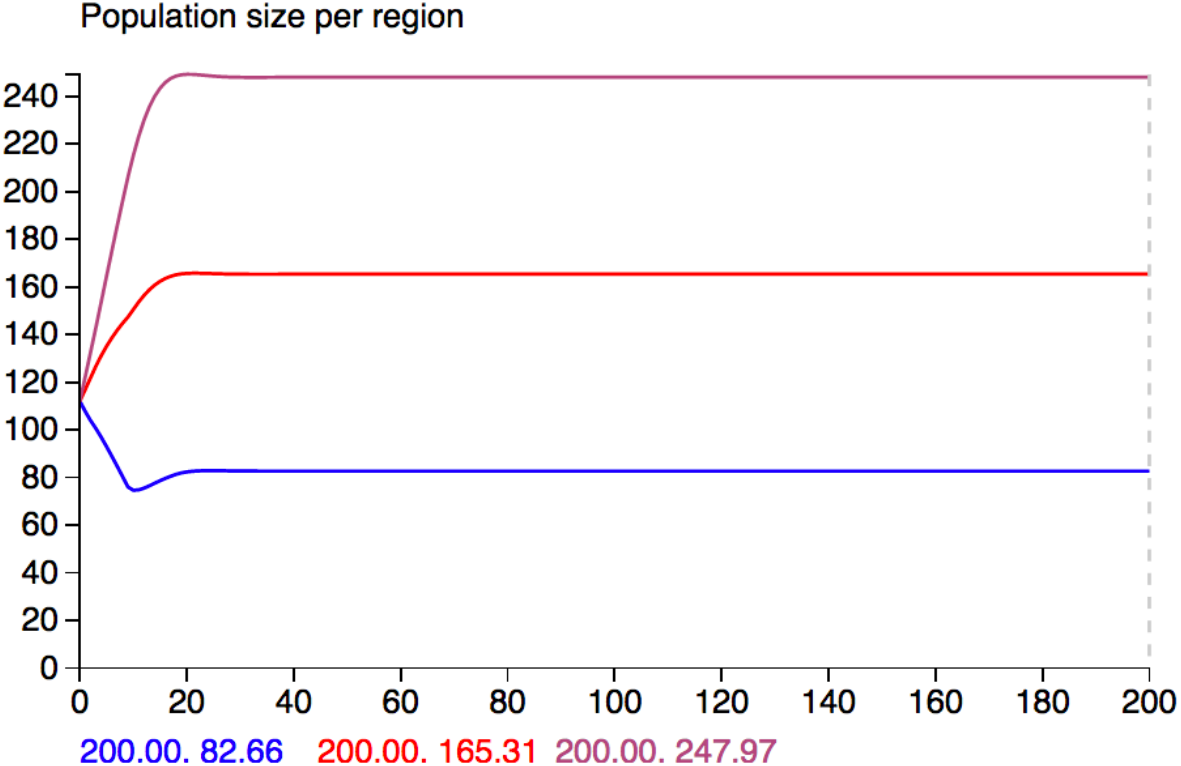
Carrying capacity parameters *c*=50, 100, and 150 produce equilibrium densities of 69, 138 and 208 individuals in Regions 1, 2, and 3 respectively. Initial values for male and female cohorts in all cases were: male=(10,9,8,7,6,5,4,3,4)´ (´ denotes the vector is transposed from a column to a row) and female==(10,9,8,7,6,5,4,3,4)´

## Appendix 3: Model Formulation and Equations

### Two-sex deterministic age-structured framework

The following is a two-sex Leslie Matrix model expressed in terms of age-sex variables (age *i*, except *n* is age *n* and older; and *f* and *m* denote female and male respectively) *x*_*if*_(*t*) and *x*_*im*_(*t*), *i* = 1,… *,n*, time *t*, and life history survival (*s_if_, s*_*im*_) and natality (*b_if_, b*_*I'm*_) parameters, where the female component of the model can be expressed as a matrix equation

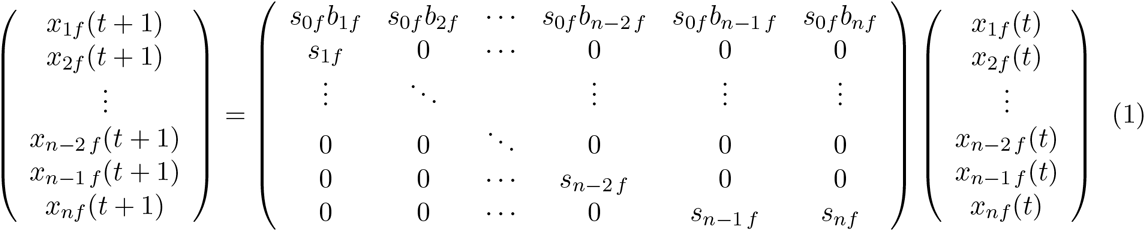
and the male component either requires doubling the dimension of the above matrix equation or augmenting the above equations with equations

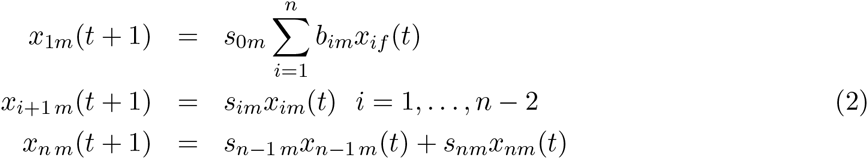
Under the assumption that males have no effect on the female component of the model, as long as there are sufficient males to fertilize sexually mature females, the growth or decline rate of the female component is determined by the eigenvalues of the matrix depicted in Equation 1. If this matrix is primitive (nonnegative and irreducible with at least one positive element) then it is known from the Perron-Frobenius Theorem that this matrix has a positive dominant eigenvalue, say λ_1_, and corresponding eigenvector, say x_1_, such the population ultimately grows (λ_1_ > 1) or declines (λ_1_ < 1) at rate λ_1_ and the solution vector x_*f*_(*t*)′ = (*x*_1*f*_(*t*),…, *x*_*nf*_(*t*)) (′ denotes vector transpose) directionally aligns with x_1_ as *t* → ∞.

Growing populations will ultimately be regulated through density-dependent mechanisms linked to resources available to each individual over each period of time. Typically, age classes likely to be most vulnerable to the effects of competition for resources are the youngest and oldest age classes, although the effects of competition on all age classes ca can be considered. A simple approach is to express the effects of density on age class *i* in terms of the ratio of an aggregated population index *B*_*il*_(*t*) and age-sex-specific available resources *R*_*il*_(*t*). Specifically, for a set of weights *w*_*ijl*_ ≥ 0, we define

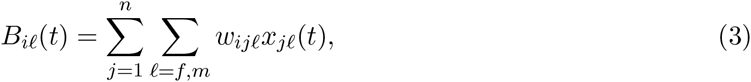
If the weights *w*_*ijf*_ and *w*_*ijm*_ are the actual or relative weights of individuals aged *j* then *B*_*i*_(*t*) is a population biomass index. The resources *R*_*i*_(*t*) can either be external inputs or systems variables that depend on the population via consumer-interaction processes. The density-dependent effects on age class *i* can be included using functions *F*_*il*_ to multiplicatively modify the values of *s*_*il*_; where, for scaling constants *c*_*il*_ > 0,

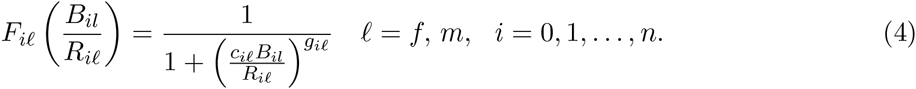
We note that 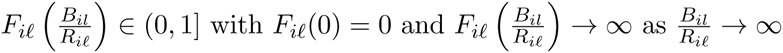.

In the simplest two-sex, density-dependent case, we assume that an age-independent birth sex-ratio variable *ρ* ∈ (0,1] (right-hand of the interval is closed since female only populations—i.e.clonal populations—are possible) applies to total birth parameters *b*_*i*_, *i* = 1,… *,n*, in which case we have

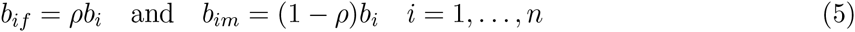
Further, if density-dependence applies only to the survival of newborns to age 1, then we have *F*_*il*_ = 0, for all *l = f, m* and *i* = 1,… *,n.* In this case, we have one density-dependent function, which we denote by

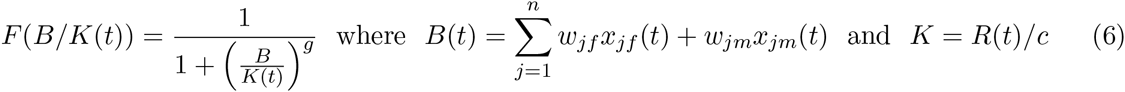
This function premultiplies only the parameters *s*_0*f*_ in Equation 1 and *s*_0*m*_ in Equation 2. (Note, in our main text, we refer to *K* as the “carrying capacity” and select the units of *c* so that the carrying capacity is scaled to adult male biomass equivalents per unit area).

**Figure.**
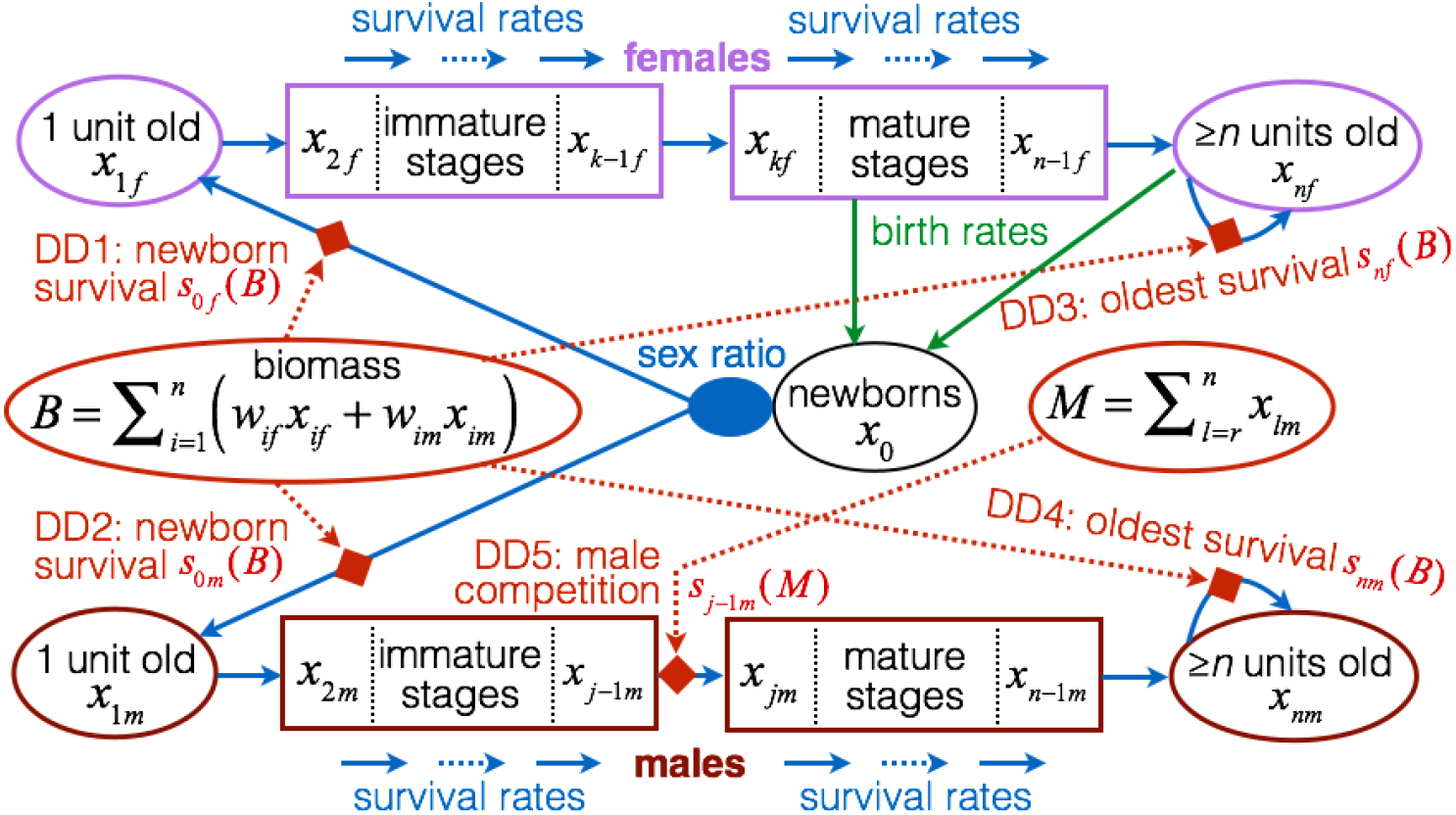
A flow diagram of a two-sex, age-structured, population model with 5 sources of density-dependent survival in the youngest (DD1, DD2) and oldest (DD3, DD4) classes that depend on the population biomass B (Eq. 3), as well as in the maturing males that depending on the number of mature males M (DD5; Eq. 4). The latter DD-survival rate depends on the number of mature males in the population, while the other DD-survival rates depend on the total biomass of the population.

### Demographic stochasticity

Demographic stochasticity arises in the context of survival of individuals when we regard survival parameters *s*_*if*_ and *s*_*im*_ as denoting probabilities that each individual survives rather than the proportion of individuals in the *i*^*th*^ age class that survive. In this case, the variables *x*_*if*_ and *x*_*im*_ are regarded as random variables *X*_*if*_ and *X*_*im*_ that are determined from binomial distributions arising from repeated Bernoulli trials of whether or not each individual survives or does not survive with probabilities *s*_*if*_ and *s*_*im*_, etc. For example, if *x*(*t*) individuals each survives the interval [*t, t* + 1) with probability *s*, then the computation of the value *x*(*t* + 1) arises from a drawing of a variable *X*(*t*) ~ BINOMIAL[*x*(*t*),*s*]. Notationally, we use the equation

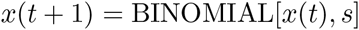
to mean that we have drawn a value from the distribution BINOMIAL[*x*(*t*), *s*] and called this value *x*(*t* + 1). Using this convention, our survival equations, incorporating the density-dependent functions given in Eqns 4, but omitting the arguments of this functions for clarity: take the form

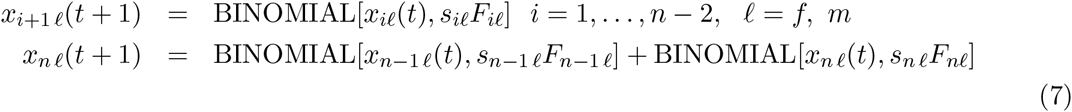
It only remains now to generate equations for *x*_1*l*_(*t* + 1), *l = f, m.* This requires that we first generate the number of newborn individuals *x*_0,*i*_(*t*) to females of age *i*, calculate the proportion of these that are female and male, using a sex-ratio probability-of-being-female parameter *ρ*_*i*_ ∈ (0,1), and then the probability that these young survive the year, where this survival may also depend on the age *i* of the mothers. To allow for this level of generality, we first calculate *x*_0,*i*_(*t*), as described below. We then calculate the total of these that may be female *x*_0,*if*_(*t*) = BINOMIAL[*x*_0_,_*i*_(*t*),*ρ*_*i*_], with the remaining *x*_*0,im*_(*t*) = *x*_0_,_*i*_(*t*) − *x*_0*,if*_(*t*) being males. Then, if s_0,*il*_ and *F*_0,*il*_ are the density independent and dependent components respectively of the survival rates of female (*l = f*) and male (*l* = *m*) young in their first year to mothers of age *i*, we finally obtain the equations:

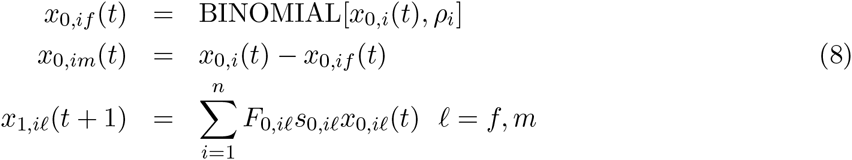

So it remains now to discuss how to generate the equation for *x*_0_(*t*). We consider the simple case where each female can have at most on young: this is the case for many large mammals, particularly herbivores (e.g. elephants, rhino, hippo, large antelope).

#### Case 1: single births

Each individual female in age class *i* will give birth to 1 or 0 individuals with probabilities *b*_*i*_ ∈ (0,1) and (1 − *b*_*i*_). In this case, if there are *x*_*i*_, _*f*_(*t*) females at time *t* then

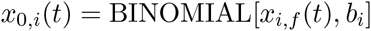

#### Case 2: binomial with maximum number of multiple births

Each individual can have at most 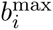 (which must be an integer), with the expected number of young being 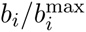, with the actual litter size being binomial. In this case we have *x*_*i*,*f*_(*t*) drawings from the distribution 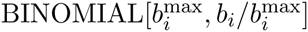 to obtain

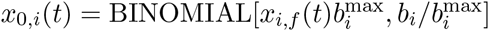
which generalizes the case above where we have *b*_*i*_^*max*^ = 1.

### Environmental stochasticity

Environmental stochasticity is easy to incorporate into the model in two different ways. First, in the functions in Eqns. 4, the resources *R*_*il*_(*t*) available to individuals in demographic class *il* can vary stochastically from one time period to the next. This is particularly easy to characterize if *R*_*il*_(*t*) is treated purely as an input rather than a systems variable interacting with its consumers. Second, survival probabilities themselves can be treated as stochastic variables rather than constants. This can be done both in terms of large infrequent perturbations due to epidemics or other types of environmental catastrophes. Extractions due to predation or human activities, however, are considered elsewhere.

A particularly simple environmental variable approach to including stochasticity, which is the one we took in developing our Nova web app, is to select a parameter γ ∈ [0,1] such that γ = 0 corresponds to the absence of environmental stochasticity and γ = 1 maximum stochasticity. We then flip a coin and decide if it is heads we select a female survival value at random on [*s*_0*f*_, *s*_0*f*_ + γ(1 − *s*_0*f*_)] to obtain 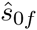. We now compute 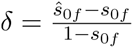 and apply the same proportional increase to male survival value to obtain 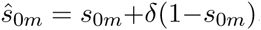. On the other hand, if the coin is tails we then select a female survival value at random on [*s*_0*f*_ – γ*s*_0*f*_, *s*_0*f*_]. We now compute 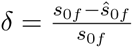 and apply the same proportional decrease to male survival value to obtain 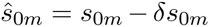. Note that this approach implies that expected survival is not equal to the nominal survival s0 as the stochasticity increases, unless *s*_0_ = 0.5: if *s*_0_ > 0.5 then the expected survival will be less than the nominal survival (since *s*_0_ < 1 − *s*_0_) and vice versa if *s*_0_ < 0.5.

Environmental variation through dependency of resources *R*_*il*_(*t*) can be incorporated using a time-series model. In particular, a relatively simple model (Murdoch et al Nature 2002) that applies to a collection of age-sex-classes denote by I (e.g. all adults), with *B*_*I*_(*t*) defined in terms of a weighted sum of individuals over the index set *I*, *F*_*I*_ (*B*_*Il*_, *R*_*Il*_) a function of the form defined in Eqn 4, and parameters *c*_*I*_ > 0, 0 < *d*_*I*_ < 1 and λ_*I*_:

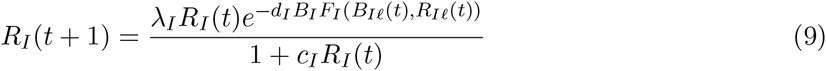
where the parameter *c*_*I*_ can be treated as a random variable (i.e. this essentially treats the carrying capacity of *R*_*I*_ as a random variable) belonging to an appropriate distribution defined on (0, ∞); for example, a lognormal distribution:

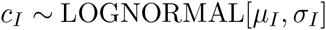
Whatever distribution is used, if c_*I*_ and *B*_*I*_ are small then the resource grows at a per capita rate λ_*I*_, while large c_*I*_ or intermediate *B*_*I*_ reduce this growth rate (note: if *g*_*I*_ > 1, which it invariably is as discussed in Getz 1996, then very large *B*_*I*_ leads to very low survival and a collapse in the population, which to some extent mitigates against resource devastation).

In the case of simply treating the parameter *K*_*I*_ in Equation 6 as a random variable

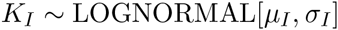
if the mean and standard deviation of the carry-capacity of input values 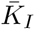 and *S*_*I*_ respectively, then it is known that the values of *µ*_*I*_ and *σ*_I_ in the LOGNORMAL distribution used to generate values *K*_*I*_(*t*) for each interval *t* are

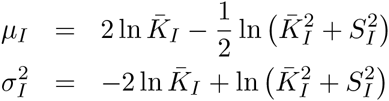

### Metapopulations and migration

If the population has a metapopulation structure, then the simplest approach to capturing this structure is to treat the subpopulations in each of *h* areas as homogeneous entities linked by movements of individuals among subpopulations. Let the *j*^*th*^ of these subpopulations be represented by a vector (note we are using ′ to denote transpose that allows as to list vectors in row form rather then column form)

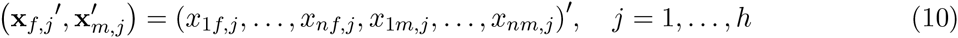
Movement of individuals among areas can then be scheduled after demographic (births, survival and aging), harvesting and stocking rates have been accounted for in each subpopulation. This will be handled using a Markov transition matrix approach in which each individual of age-sex class (*il*) has a current integer state value *η* and a next integer state value *ζ*, where *η, ζ* = 1,…, *h* designate the subpopulations of *origination (*η) and destination (*ζ*).

For each age-sex class (*il*), we create a stochastic movement transition matrix *M*^(*il*)^ with elements 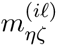 by definition satisfying 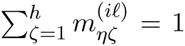 Thus, if there are 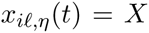 individuals of age-sex class (*il*) in subpopulation *η* then the distribution of these individuals in the different subpopulations at time *t* + 1 is given by 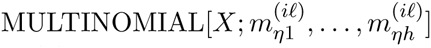. All the remains now is to compute the values in the matrix *M*^(*il*)^ over the range *i* = 1, …,*n* and *l = m*, *f*. In many case the matrices may not vary across various subranges, such as all adult females, and so on. Additionally, in many case, only certain age-sex classes may move, such as males at first age for maturity; in which case all the elements of the remaining matrices are 0.

Consider an overall movement processes that is a concatenation of the following three components:

1. an age-sex class movement propensity *q*_*il*_ that is independent of location state *η*
2. a connectivity matrix *C* of elements 0 ≤ *c*_*ηζ*_ ≤ 1 that determines how relatively easy it is for individuals to move between any two locations *η* and ζ. Thus for example, if *c*_32_ = 0 then individuals cannot move between locations 3 and 2, though *c*_23_ > 0 would imply that they can still move between locations 2 and 3. In the most general case, *C* could depend on age-sex class, since individuals in different classes may have different movement capabilities. But we will not consider this level of generality here.
3. a subpopulation attractivity vector *α*, with elements *α*_ζ_, that is dependent only on the relative attractivity of the different subpopulations (such as the values of the youngest age-class density-dependent factors *F*_0,*ζ*_(*t*) for the subpopulations, should they exist).
Once all these values have been entered or determined for each class (*il*) of individuals that will move, then we can calculate the movement matrix entries as

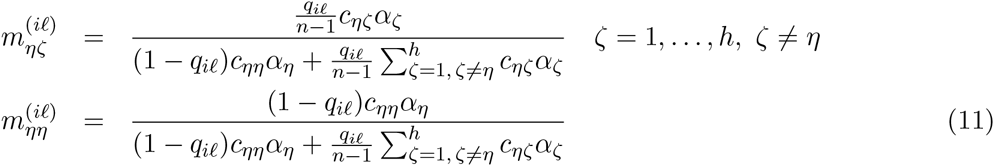
noting that, in effect, the *η*^*th*^ entry of the computation 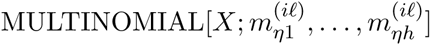 represents those individuals who either did not leave subpopulation *η* in the first place, or went on a walk-about and then returned back to their originating population after sampling the attractivity of the other populations: either interpretation holds.

#### Notes on Code Implementation

The vectors 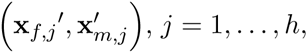 (each of dimension 2*n* that are used in the movement algorithm (cf. Equation 10 above) represent the numbers in each of the 2*n* (*il*)-age-sex classes for each of the *j* = 1, …,*h* subpopulations after computations at the lower level have been carried out with respect to survival, extraction (harvesting), and stocking. Before these vectors are then passed back for demographic updating (calculation of the age transitions *x*_*i*+1*l*_(*t*+1) in each of the *h* subpopulations), we calculate movement using the values 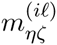 computed in equations above in the *2n* × *h* computations

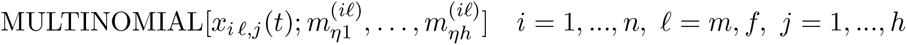

### Extraction and exploitation

Unless predators are explicitly identified, survival estimates from life tables usually implicitly include losses to predation at static background rates. If predators need to be explicitly included as interacting dynamical with the population of interest, which will be the case if the population is the primary food source for a population of predators, then such descriptions can be included. The details, however, will differ depending on the type of prey-predator interaction considered. In a number of conservation biology problems, interest exists in the fate of populations subject to extraction by humans, either because of legal harvesting or because of illegal pouching (e.g. rhinos). In each case, the state of the population after demography has been accounted for over each time interval, additional individuals can be removed using appropriate deterministic or stochastic rules to simulated the effects of exploitation by humans (as discussed in the main text).

### Pseudo-extinction statistics

Populations are typically considered to be extinct in the wild, once the last remaining individuals in a natural area have been removed to sanctuaries for protection and breeding programs, where the latter may later be used to restock natural areas under so-called ‘reintroduction programs’. Population levels at which such interventions occur are called pseudo-extinction levels. A pseudo extinction criterion is thus a combination of adult female and male levels (treated separately or combined) at which the breeding population drops below a critical level (for combined: < *X*_*cc*_) or the number of mature females or males drops below critical levels on a specified interval of time, say [0,*T*].

## References

Armstrong, D. P. and P. J. Seddon. 2008. Directions in reintroduction biology. Trends in Ecology & Evolution 23:20–25.

Barnosky, A. D., N. Matzke, S. Tomiya, G. O. U. Wogan, B. Swartz, T. B. Quental, C. Marshall, J. L. McGuire, E. L. Lindsey, K. C. Maguire, B. Mersey, and E. A. Ferrer. 2011. Has the Earth's sixth mass extinction already arrived? Nature 471:51–57.

Beissinger, S. R. and M. I. Westphal. 1998. On the use of demographic models of population viability in endangered species management. Journal of Wildlife Management:821–841.

Beverton, R. J. H. and S. J. Holt. 1957. On the dynamics of exploited fish populations. Ministry of Agriculture and Fisheries.

Brito, D and G. A. B. Da Fonseca. 2006. Evaluation of minimum viable population size and conservation status of the long-furred woolly mouse opossum Micoureus paraguayanus: An endemic marsupial of the Atlantic Forest. Biodiversity and Conservation 15:1713–1728.

Brodie, J. F., J. Muntifering, M. Hearn, B. Loutit, R. Loutit, B. Brell, S. Uri-Khob, N. Leader-Williams, and P. du Preez. 2011. Population recovery of black rhinoceros in north-west Namibia following poaching. Animal Conservation 14:354–362.

Caswell, H. 2001. Matrix population models : construction, analysis, and interpretation. 2nd edition. Sinauer Associates, Sunderland, Mass.

Chapron, G., D. G. Miquelle, A. Lambert, J. M. Goodrich, S. Legendre, and J. Clobert. 2008. The impact on tigers of poaching versus prey depletion. Journal of Applied Ecology 45:1667–1674.

Conway, A. J. and P. S. Goodman. 1989. Population Characteristics and Management of Black Rhinoceros Diceros-Bicornis-Minor and White Rhinoceros Ceratotherium-Simum-Simum in Ndumu-Game-Reserve, South-Africa. Biological Conservation 47:109–122.

Crone, E. E., E. S. Menges, M. M. Ellis, T. Bell, P. Bierzychudek, J. Ehrlen, T. N. Kaye, T. M. Knight, P. Lesica, W. F. Morris, G. Oostermeijer, P. F. Quintana-Ascencio, A. Stanley, T. Ticktin, T. Valverde, and J. L. Williams. 2011. How do plant ecologists use matrix population models? Ecology Letters 14:1–8.

Desharnais, R. A., R. F. Costantino, J. M. Cushing, S. M. Henson, B. Dennis, and A. A. King. 2006. Experimental support of the scaling rule for demographic stochasticity. Ecology Letters 9:537–547.

Easterling, M. R., S. P. Ellner, and P. M. Dixon. 2000. Size-specific sensitivity: Applying a new structured population model. Ecology 81:694–708.

Ellis, E. C. 2011. Anthropogenic transformation of the terrestrial biosphere. Philosophical Transactions of the Royal Society a-Mathematical Physical and Engineering Sciences 369:1010–1035.

Ellner, S. P. and M. Rees. 2006. Integral projection models for species with complex demography. American Naturalist 167:410–428.

Engen, S., R. Lande, B.-E. Sæther, and H. Weimerskirch. 2005. Extinction in relation to demographic and environmental stochasticity in age-structured models. Mathematical Biosciences 195:210–227.

Fieberg, J. and S. P. Ellner. 2001. Stochastic matrix models for conservation and management: a comparative review of methods. Ecology Letters 4:244–266.

Foley, J. A., N. Ramankutty, K. A. Brauman, E. S. Cassidy, J. S. Gerber, M. Johnston, N. D. Mueller, C. O'Connell, D. K. Ray, P. C. West, C. Balzer, E. M. Bennett, S. R. Carpenter, J. Hill, C. Monfreda, S. Polasky, J. Rockstrom, J. Sheehan, S. Siebert, D. Tilman, and D. P. M. Zaks. 2011. Solutions for a cultivated planet. Nature 478:337–342.

Gaillard, J-M., M. Festa-Bianchet, and N. G. Yoccoz. 1998. Population dynamics of large herbivores: variable recruitment with constant adult survival. Trends in Ecology & Evolution 13:58–63.

Getz, W. M. 2011. Biomass transformation webs provide a unified approach to consumer– resource modelling. Ecology Letters 14:113–124.

Getz, W. M. and R. G. Haight. 1989. Population Harvesting: Demographic Models of Fish, Forest, and Animal Resources. (MPB-27). princeton university press.

Getz, W. M., R. M. Salter, and N. Sippl-Swezey. 2015. Using Nova to construct agent-based models for epidemiological teaching and research.*in* Proceedings of the 2015 Winter Simulation Conference.

Godfray, H. C. J., J. R. Beddington, I. R. Crute, L. Haddad, D. Lawrence, J. F. Muir, J. Pretty, S. Robinson, S. M. Thomas, and C. Toulmin. 2010. Food Security: The Challenge of Feeding 9 Billion People. Science 327:812–818.

Heppell, S. S. 1998. Application of life-history theory and population model analysis to turtle conservation. Copeia:367–375.

Kettenring, K. M., B. T. Martinez, A. M. Starfield, and W. M. Getz. 2006. Good practices for sharing ecological models. Bioscience 56:59–64.

Lacy, R and J. Pollak. 2012. Vortex: A stochastic simulation of the extinction process. Version 10.0. Chicago Zoological Society, Brookfield, Illinois.

Lacy, R. C. 1993. VORTEX: a computer simulation model for population viability analysis. Wildlife Research 20:45–65.

Lacy, R. C. 2000. Structure of the VORTEX simulation model for population viability analysis. ecological Bulletins:191–203.

Law, P. R., B. Fike, and P. C. Lent. 2014. Birth sex in an expanding black rhinoceros (Diceros bicornis minor) population. Journal of Mammalogy 95:349–356.

Linklater, W. L., K. Adcock, P. du Preez, R. R. Swaisgood, P. R. Law, M. H. Knight, J. V. Gedir, and G. I. H. Kerley. 2011. Guidelines for large herbivore translocation simplified: black rhinoceros case study. Journal of Applied Ecology 48:493–502.

Marker, L. and C. C. Fund. 2012. International cheetah (Acinonyx jubatus) studbook. Cheetah Conservation Fund, Namibia.

Menges, E. S. 2000. Population viability analyses in plants: challenges and opportunities. Trends in Ecology & Evolution 15:51–56.

Merow, C., J. P. Dahlgren, C. J. E. Metcalf, D. Z. Childs, M. E. Evans, E. Jongejans, S. Record, M. Rees, R. Salguero - Gómez, and S. M. McMahon. 2014a. Advancing population ecology with integral projection models: a practical guide. Methods in Ecology and Evolution 5:99–110.

Merow, C., A. M. Latimer, A. M. Wilson, S. M. McMahon, A. G. Rebelo, and J. A. Silander. 2014b. On using integral projection models to generate demographically driven predictions of species' distributions: development and validation using sparse data. Ecography 37:1167–1183.

Metzger, K. L., A. R. E. Sinclair, R. Hilborn, J. G. C. Hopcraft, and S. A. R. Mduma. 2010. Evaluating the protection of wildlife in parks: the case of African buffalo in Serengeti. Biodiversity and Conservation 19:3431–3444.

Minteer, B. A. and J. P. Collins. 2010. Move it or lose it? The ecological ethics of relocating species under climate change. Ecological Applications 20:1801–1804.

Moffitt, E. A., J. W. White, and L. W. Botsford. 2011. The utility and limitations of size and spacing guidelines for designing marine protected area (MPA) networks. Biological Conservation 144:306–318.

Morris, W. F. and D. F. Doak. 2002. Quantitative conservation biology : theory and practice of population viability analysis. Sinauer Associates, Sunderland, Mass.

Mullon, C., P. Freon, and P. Cury. 2005. The dynamics of collapse in world fisheries. Fish and Fisheries 6:111–120.

Ostrom, E., J. Burger, C. B. Field, R. B. Norgaard, and D. Policansky. 1999. Sustainability -Revisiting the commons: Local lessons, global challenges. Science 284:278–282.

Quinn, T. J. and R. B. Deriso. 1999. Quantitative fish dynamics. Oxford University Press, New York.

Rees, M., D. Z. Childs, and S. P. Ellner. 2014. Building integral projection models: a user's guide. Journal of Animal Ecology 83:528–545.

Salter, R. M. 2013. Nova: A modern platform for system dynamics, spatial, and agent-based modeling. Procedia Computer Science 18:1784–1793.

Seddon, P. J., C. J. Griffiths, P. S. Soorae, and D. P. Armstrong. 2014. Reversing defaunation: Restoring species in a changing world. Science 345:406–412.

Urban, M. C. 2015. Accelerating extinction risk from climate change. Science 348:571–573.

Watson, J. E. M., N. Dudley, D. B. Segan, and M. Hockings. 2014. The performance and potential of protected areas. Nature 515:67–73.

Weinbaum, K. Z., J. S. Brashares, C. D. Golden, and W. M. Getz. 2013. Searching for sustainability: are assessments of wildlife harvests behind the times? Ecology Letters 16:99–111.

Wilmers, C. C. and W. M. Getz. 2004a. Simulating the effects of wolf-elk population dynamics on resource flow to scavengers. Ecological Modelling 177:193–208.

Wilmers, C. C. and W. M. Getz. 2004b. Simulating the effects of wolf-elk population dynamics on resource flow to scavengers. Ecological Modelling 177:193–208.

Wisdom, M. J., L. S. Mills, and D. F. Doak. 2000. Life stage simulation analysis: Estimating vital-rate effects on population growth for conservation. Ecology 81:628–641.

Wittemyer, G. 2011. Effects of Economic Downturns on Mortality of Wild African Elephants. Conservation Biology 25:1002–1009.

